# Semi-automated background removal limits loss of data and normalises the images for downstream analysis of imaging mass cytometry data

**DOI:** 10.1101/2020.11.26.399717

**Authors:** Marieke E. Ijsselsteijn, Antonios Somarakis, Boudewijn P.F. Lelieveldt, Thomas Hollt, Noel F.C.C. de Miranda

## Abstract

Imaging mass cytometry (IMC) allows the detection of multiple antigens (approximately 40 markers) combined with spatial information, making it a unique tool for the evaluation of complex biological systems. Due to its widespread availability and retained tissue morphology, formalin-fixed, paraffin-embedded (FFPE) tissues are often a material of choice for IMC studies. However, antibody performance and signal-to-noise ratio can differ considerably between FFPE tissues as a consequence of variations in tissue processing, including fixation. We investigated the effect of immunodetection-related signal intensity fluctuations on IMC analysis and phenotype identification in a cohort of twelve colorectal cancer tissues. Furthermore, we explored different normalisation strategies and propose a workflow to normalise IMC data by semi-automated background removal, using publicly available tools. This workflow can be directly applied to previously obtained datasets and considerably improves the quality of IMC data, thereby supporting the analysis and comparison of multiple samples.

## Introduction

Mass cytometry has advanced as an important technology for the characterisation of cellular contextures in health and disease (1-6). A major advantage of mass cytometry is its ability to simultaneously acquire over 40 markers. The high-level of multiplexing is made possible via the use of antibodies conjugated to heavy metal isotopes rather than fluorescent tags (7). Cells are labelled with these and led into a CyTOF (Cytometry by time-of-flight) instrument, where heavy metal abundance is measured, per cell, by time-of-flight mass spectrometry (8). Technological advancements in the field have made it possible to image tissue sections as opposed to single cells, allowing for the preservation of spatial information (9). Imaging mass cytometry (IMC) allows the analysis of, among others, archival tissue samples in the form of formalin-fixed paraffin-embedded (FFPE) or fresh frozen (FF) tissue. Tissue sections are labelled with metal-conjugated antibodies and ablated in small portions (typically 1 µm^2^ = 1 pixel). The ablated tissue is then analysed with the CyTOF instrument. The pixel data is processed into an image, thereby allowing the visualization of phenotypes and incorporation of spatial information in subsequent analyses.

IMC users have already contributed with a number of studies aimed at optimising the use of this technology, including: a strategy to address signal spill-over during heavy metal detection (10) as well as methodologies to aid the implementation of large antibody panels for FFPE (11) or FF (12) tissues. Schulz and colleagues demonstrated the potential of combining protein and RNA *in situ* detection with IMC (13). Furthermore, IMC has been used to comprehensively study tissue architectures and cellular composition of breast cancers (14) and pancreatic tissues affected by type 1 diabetes (15, 16), among other applications. The increasingly widespread application of IMC for the characterisation of tissues is accompanied by the need to develop analytical tools that can handle large and complex datasets where, for instance, signal to noise ratio fluctuates across samples. The general pipeline for IMC analysis involves the creation of cell segmentation masks with ilastik (17) and CellProfiler (18), after which the resulting image-stacks and masks are processed by dedicated software packages like HistoCAT (19) or ImaCytE (20).

The majority of current IMC studies make use of FFPE tissues, due to their widespread availability in tissue archives, good morphology after fixation, and the possibility to build tissue microarrays. For the interpretation of immunohistochemistry data on FFPE tissues, it has long been recognised that antibody performance and signal detection can vary considerably between specimens. This can be explained by the use of different fixation times, size of tissue during fixation, dehydration of the tissue after fixation, the age of the FFPE tissue block or how long the tissue slides have been stored before immunodetection (21-24). Moreover, particularly impactful and difficult to control, is the ischemia period that concerns the time between the collection of a tissue and its fixation. Ischemia can cause a number of artefacts due to autolysis, protein degradation, or the drying of the outer layer of the tissue (22-26). Therefore, the comparison of intensities of antibody signal between different FFPE tissues is not general practice in the evaluation of immunohistochemistry results.

In this work we investigated three methods for the processing of IMC data. We first analysed an IMC dataset without normalisation and compared this to two normalisation strategies: background removal using manual thresholding or a semi-automated method. Both approaches were followed by per-pixel binarization of marker intensity. After comparing the three approaches we propose a workflow for the analysis of tissues that makes use of publicly available tools to generate processed IMC data. Importantly, we implemented a normalisation strategy that overcomes immunodetection intensity variations across samples and considerably improves the quality of IMC data.

## Methodology

### Tissue Material

FFPE blocks from 12 colorectal cancers were obtained from the department of Pathology of the Leiden University Medical Centre (Leiden, The Netherlands). Samples were anonymised and handled according to the medical ethical guidelines described in the Code of Conduct for Proper Secondary Use of Human Tissue of the Dutch Federation of Biomedical Scientific Societies. Colorectal cancer tissues were cut into 4 μm sections and placed on silane-coated glass slides (VWR, Radnor, PA, USA).

### Imaging mass cytometry immunodetection and acquisition

Antibodies employed in this study were conjugated to purified lanthanide metals using the Maxpar antibody labelling kit and protocol (Fluidigm, San Francisco, CA, USA). Antibodies were eluted in 50 µl antibody stabilizer solution (Candor Bioscience, Wangen im Allgäu, Germany) supplemented with 0,05 % sodium azide and 50 µl W-buffer (Fluidigm). After conjugation, all antibodies were tested by IHC on 4 µm tonsil tissue to confirm that the labelling process did not affect antibody performance. IMC immunodetection was performed following the methodology published previously by our lab (11) using the antibodies and conditions described in supplementary table 1. Tissue sections were ablated within a week after immunodetection by using the Hyperion mass cytometry imaging system (Fluidigm). The Hyperion was autotuned using a 3-element tuning slide (Fluidigm) as described in the Hyperion imaging system user guide. An extra minimum threshold of 1500 mean duals of ^175^Lu was used. Four 1000×1000 µm regions of interest per sample were selected based on haematoxylin and eosin (H&E) stains on consecutive slides and ablated at 200 Hz. Data was exported as .MCD files and .txt files and visualised using the Fluidigm MCD viewer. For downstream analysis, the .MCD files were transformed to either 32-bit multi-tiff or single-marker tiff images in the MCD viewer.

### Creation of single cell masks

For each sample, one tiff image was exported from the MCD viewer, combining the keratin and vimentin expression as well as DNA detection. ilastik ((17) v1.3.3) was used to create masks for nuclei (based on the DNA signal), cytoplasm/membrane (based on keratin and vimentin expression) and background (based on the absence of signal in the DNA, keratin and vimentin image). ilastik’s random forest classifier was trained using manually assigned pixels that underwent Gaussian smoothing (ilastik feature settings: 0.3, 0.7 and 1.0 sigma for colour/intensity, edge and texture). Training was performed on 12 images (one representative image per sample) after which the classifier was applied to all images in the dataset and data was exported as probability maps indicating the likelihood of each pixel corresponding to nucleus, cytoplasm/membrane or background. In CellProfiler ((18) v2.2.0) the probability maps were used to create single cell masks for all samples. All masks were compared to the original IMC images to validate the cell segmentation procedure.

### Background removal and data normalisation

To address variations in immunodetection signal intensity between samples, two normalisation approaches were applied: 1) manual thresholding to remove background noise in MCD viewer or 2) semi-automated background removal in ilastik, both followed by binarization of pixel values.

1. Manual thresholding was done using the MCD viewer by inspecting each marker and setting a minimum intensity/mean duals threshold to remove background noise. The threshold was identified by visual inspection, based on the user’s knowledge of the expected immunodetection pattern and corresponding IHCs of the protein in question. After setting a threshold for each marker, the data was saved as .txt files containing all previously defined thresholds. This process was repeated for all images. The threshold .txt files were loaded into ImaCytE with the, in the MCD viewer, created multi-tiff images and cell segmentation masks. In ImaCytE, the thresholds were applied to the images and pixel intensity values were binarized (i.e., all pixels below the threshold were set to 0 and all pixels above threshold were set to 1). Normalised cell intensities were then defined as the frequency of positive pixels, per cell.
2. Semi-automated background removal was done on single marker tiff images, exported from MCD viewer. The images corresponding to a single marker across the entire cohort were loaded into ilastik and a small amount (i.e., approximately 1%) of pixels were assigned to either ‘signal’ or ‘background’ in 12 images (one representative image per sample). To facilitate the pixel annotation, the images were first sharpened in MATLAB. More specifically, outliers were removed through saturation of all pixels with values lower than the 1^st^ and higher than the 99^th^ percentile. Then, after Gaussian smoothing (ilastik feature settings: 0.7, 1.0 and 1.6 sigma for colour/intensity, edge and texture), the random forest classifier automatically classified signal and background pixels. After training on at least 12 images (1 for each sample), the classifier was applied to all images in the dataset and the data was exported as binary expression maps with the ‘background’ pixels set to 0 and the ‘signal’ pixels set to 1. This approach was repeated for each marker. A folder was created for each image containing the binary expression maps of all markers and the previously created cell segmentation masks. These were loaded into ImaCytE and normalised marker expression was defined by the relative frequency of positive pixels, per cell.

### Single cell clustering and phenotype calling

Marker expression per cell was obtained by processing cell segmentation masks with their corresponding pixel intensity files in ImaCytE, for the generation of FCS files. For the analyses without normalisation or with manual thresholding, multi-tiff images and their corresponding cell segmentation masks were employed. For the analysis with semi-automated normalisation, segmentation files were loaded together with binary expression maps. Single-cell FCS files containing mean pixel values per cell (non-normalised) or relative frequency of positive pixels (normalised) were then exported from ImaCytE and analysed by t-SNE ((27) t-distributed stochastic neighbourhood embedding) in Cytosplore. (28) Cells forming visual neighbourhoods in the t-SNE embedding were grouped using Mean-shift clustering and exported as separate FCS files. The resulting subsets were imported back in ImaCytE for visualisation of the subsets in the segmentation masks. All subsets were visually inspected by comparing their localisation in the segmentation masks to the original images.

## Results

### Variation in signal to noise ratio between FFPE samples influences unsupervised image analysis

Immunodetection in FFPE tissues is complicated by variations in antigen availability and accessibility across samples due to tissue processing and fixation procedures. To understand its implications to the quality of IMC data, we analysed 48 images generated from 12 colorectal cancer samples. FFPE tissues were labelled with a 36 antibody panel and four regions of interest (ROI) of 1 mm^2^ were ablated, per tumour, by the Hyperion imaging mass cytometer. Visual inspection of the images using the MCD viewer showed that large differences exist in immunodetection intensity of the same antigen between tissues, which cannot be always explained by biological variation (Figure 1a). To determine the impact of these fluctuations on downstream analyses, the added-value of two normalisation approaches was investigated in comparison to the IMC analysis pipeline without normalisation (Figure 1b). In short, cell masks were created using ilastik and CellProfiler and were loaded into ImaCytE combined with the raw or normalised images in order to define relative antigen expression per cell. FCS files were produced, and clustering of cells was performed by t-SNE to identify cell subsets, using Cytosplore. Next, the phenotypes were projected back onto the cell masks in ImaCytE for visualisation and spatial analysis.

**Figure 1.**
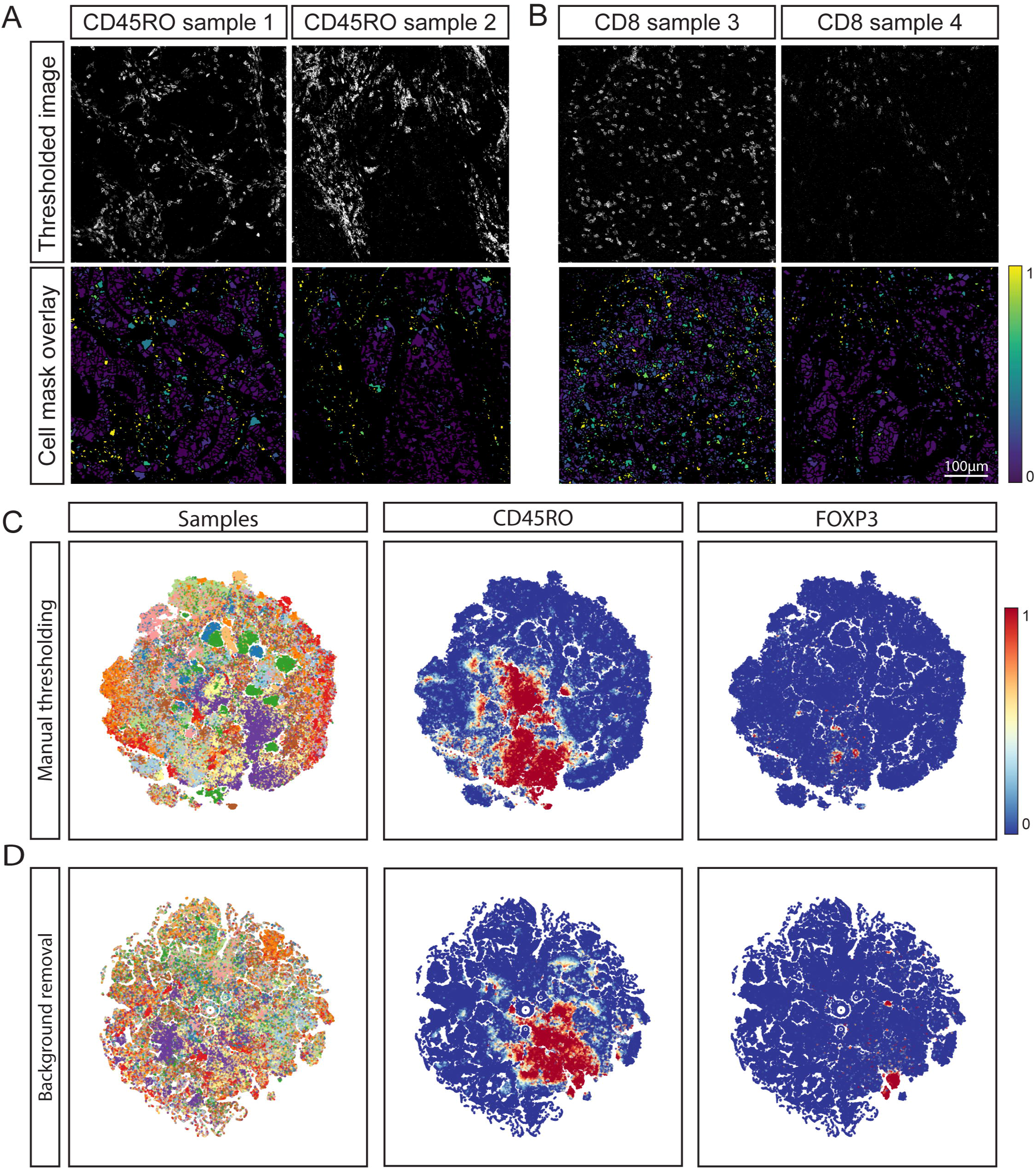
(a) CD163 expression pattern in three samples. A signal range of 3-20 mean duals was set for all images in the MCD viewer, but differences in signal intensity and noise are observed between images. (b) Workflow for IMC FFPE imaging and data processing, including the three tested data processing approaches where either no normalisation was performed, binarization by manual thresholding or binarization by semi-automated background removal.

First, we visualised immunodetection signal intensities, without normalisation, per antibody, on all cell mask overlays where antibody signal was displayed as mean pixel intensity (Figure 2a and 2b, lower panel). Where differences were observed, the original IMC images were inspected, showing that fluctuations in intensity on the cell masks generally corresponded to variations in signal-to-noise ratios. This resulted in either overestimation (Figure 2a) or underestimation (Figure 2b) of cells positive for a marker with variable signal-to-noise between samples. Furthermore, similar immune cell subsets could be assigned to distinct immune cell populations as a consequence of high variability in signal intensity for some markers between samples.

**Figure 2.**
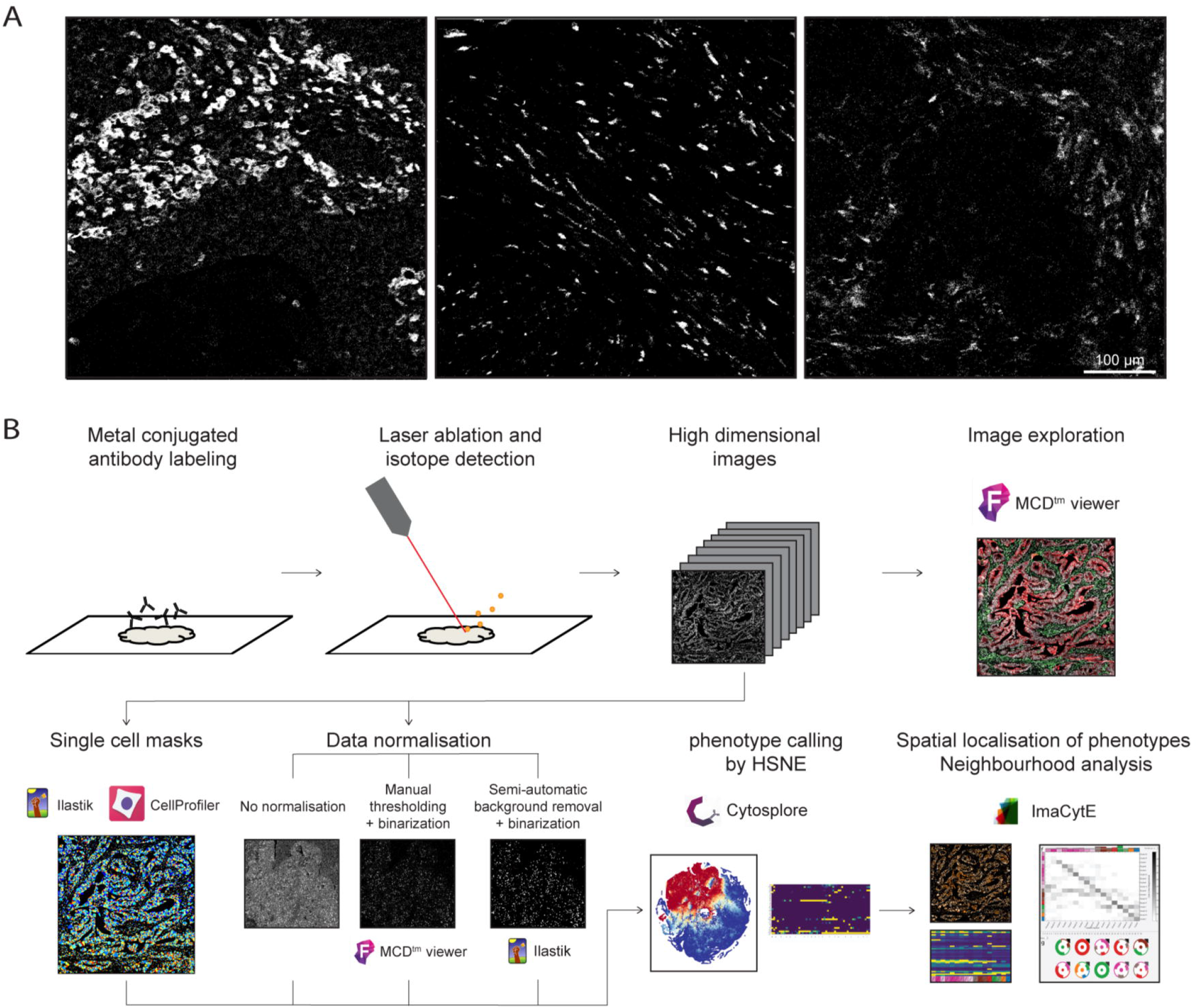
(a) correlation between original MCD image and cell expression after mask overlay. Variation between images occurs due to differences in background as seen for CD45RO between sample 1 and 2 and (b) variation in signal intensity as observed for CD8 between sample 3 and 4. Signal in the cell mask ranged from 0 – 10 mean duals. (c) t-SNE analysis embedding of single cell data from all 48 images in which cells were coloured by sample of origin in the -t-SNE embedding. CD45RO and CD8 examplify the spread of positive cells throughout the t-SNE embedding.

To test this, a t-SNE embedding was computed using the single cell maker expression data extracted from 48 images. The embedding contained 393727 cells and was then visualised in a two-dimensional scatterplot with sample IDs and expression of each marker shown by colour coding. It was observed that cells belonging to the same sample were localised in close proximity suggesting sample-inherent bias (Figure 2c, Supplementary Figure 1, 2a). Furthermore, cells with a similar marker profile were scattered throughout the t-SNE embedding rather than clustering together, suggesting that the intensity range and signal-to-noise variation between samples overshadowed cell type differences and similarities. Finally, cells positive for FOXP3, CD20 or CD103, markers with a low signal-to-noise ratio, did not cluster in the t-SNE analysis (Supplementary Figure 1).

### Manual thresholding and binarization normalises IMC inter-sample variation for automated downstream analysis

A methodology was devised to test whether the observed immunodetection variation could be overcome by normalising the IMC data, while minimising data loss, for downstream analysis. This approach utilised a user-defined minimum signal threshold for each marker and then pixel binarization of the dataset. To confirm whether this approach was sufficient for reliable downstream analysis, we visualised the normalised marker expression on the cell masks. Indeed, setting a minimum signal threshold resulted in comparable marker detection between samples (Figure 3a and 3b) However, for several markers, it was observed that the number of positive cells in the cell mask was considerably lower than in the original images, suggesting that setting a manual threshold to remove noise also leads to loss of true positives.

**Figure 3.**
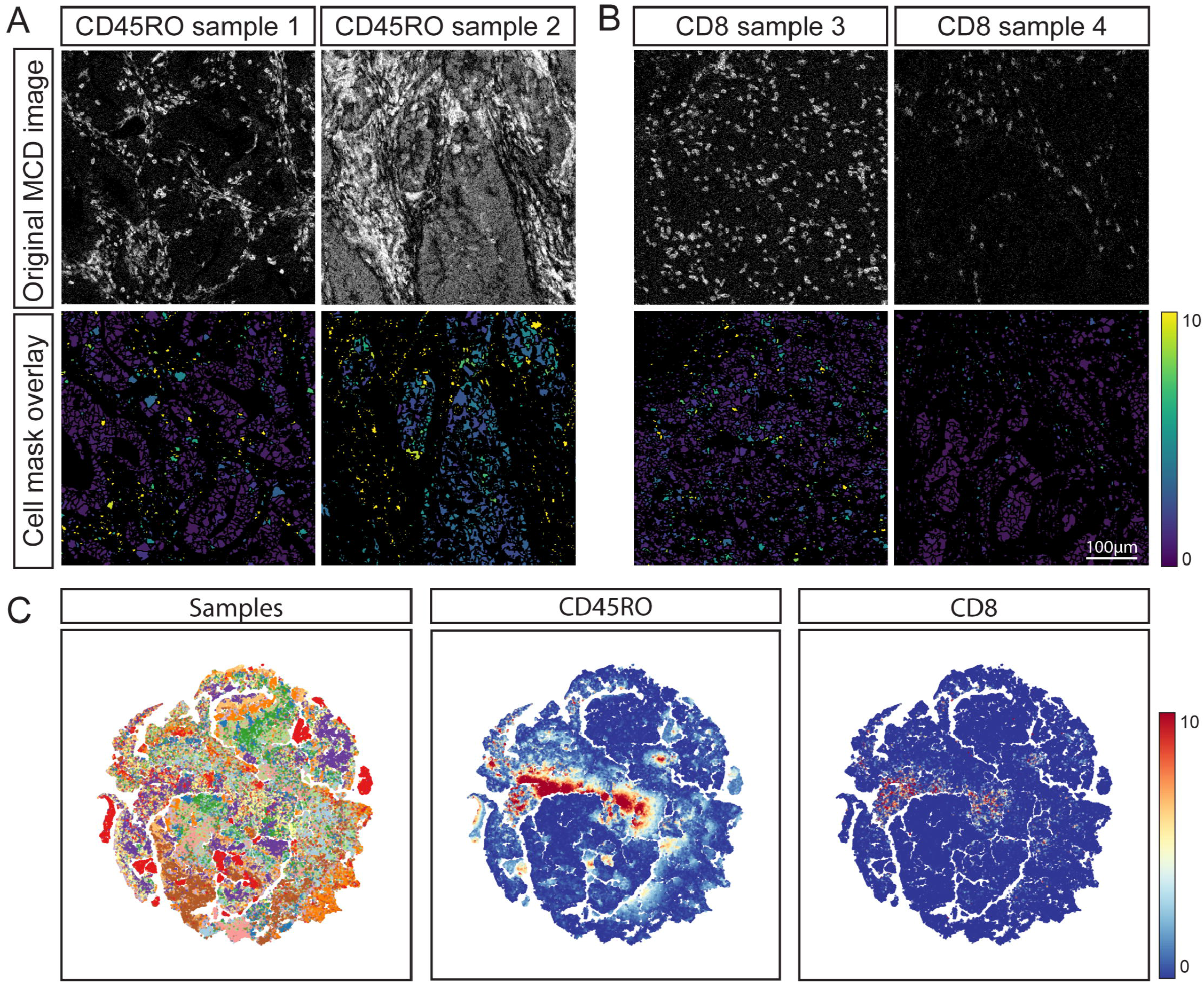
(a & b) correlation of CD45RO and CD8 between two thresholded MCD images and the mask overlay after manual thresholding and pixel binarization. Signal intensity ranges between 0 and 1 due to the visualisation as frequency of positive pixels per cell. (c) t-SNE embedding of the single cell data from all 48 images after manual thresholding and binarization. Cells were coloured by sample of origin in the t-SNE embedding. CD45RO and FOXP3 were shown to highlight the distribution of positive cells in the t-SNE embedding (d) t-SNE embedding of single cell data after semi-automated background removal and binarization. Cells were coloured by sample of origin in the t-SNE embedding.

To further assess the effect of thresholding and binarization on downstream analysis, t-SNE embedding, as described in the previous section, was performed on the single cell data obtained from the thresholded and binarized images (Figure 3c, supplementary figure 1). In contrast to the analysis of the dataset without normalisation, the majority of cells clustered based on their phenotypes but some sample-specific clustering remained (Figure 3c, Supplementary figure 2b-d). Further inspection of the t-SNE embedding showed that the cells in those clusters were keratin-positive (a marker for epithelial cells) with varying combinations of HLA-DR, Ki-67 and CD15 expression, markers that are often differentially expressed between cancer cells which provides a biological basis for their sample-specific clustering. In contrast to the TSNE embedding of data without normalisation, cells with a similar marker profile clustered together (as observed for CD8, Supplementary figure 1) Furthermore, the distinction between positive and negative cells for a specific marker was more clear (as observed for CD163, Supplementary figure 1). However, in line with the observations made during visual inspection, the number of cells positive for dim markers was low compared to the original MCD images. Albeit higher than the number of positive cells observed in the analysis without normalisation. (Figures 2c, 3c and Supplementary figure 1) Thus, thresholding and binarization of pixel intensity largely resolved sample-specific clustering and allowed for comparison between samples, but does not resolve the presence of false negatives.

Although manual thresholding was found to overcome some of the challenges of analysing FFPE IMC data, its major disadvantages are that it is time consuming and subjective to errors as it requires vast knowledge of the expected labelling patterns of each marker and high inter-user variability is inevitable. Furthermore, while thresholding removes background noise, a portion of specific signal can be lost particularly when the signal-to-noise ratio is low, resulting in false-negatives. Therefore, we set out to investigate if an automated and unbiased approach could replace manual thresholding. We first visualised the pixel data in histograms for each marker to assess if pixel intensity was bimodally distributed in order to set an automatic threshold between negative and positive pixels. However, no bimodal distribution but a negative correlation between number of pixels and signal intensity was observed (Supplementary figure 3a). We then investigated the possibility to define a threshold based on intensity distribution at the cell level after applying cell masks onto the images. Initially, we set the threshold value at 1, regarding all pixels with higher intensities values as positive. Then, we visualised the data on cell level by plotting the mean of all pixels within a cell (supplementary figure 3b, g-h). Also, at cell level, no clear bimodal distribution was observed. Similarly, cut-off thresholds values between 2 and 5 resulted in similar distributions (supplementary figure 3c-f). Moreover, a threshold of 2,9 mean duals was comparable to the cut-off chosen during manual thresholding but this could not be deduced from the single-cell value distribution (supplementary figure 3d, i). Thus, an automated approach to determine a precise cut off value could not be established. Furthermore, setting a single-value threshold, as was also observed with manual thresholding, causes a trade-off between the removal of background noise and low intensity true signal and does not overcome case-specific background signal as was observed for some images and markers (e.g. CD45ro, Figure 2a).

### Semi-automated background removal limits loss of data and normalises the images for downstream analysis of IMC data

To correct for both technical noise and sample-specific background signal, a semi-automated background removal approach based on ilastik’s pixel classification algorithm was employed. For each marker, pixels corresponding to either ‘background’ or ‘signal’ were labelled and used to train a random forest classifier in ilastik. After training on 12 images per marker (or 1 image per sample) the algorithm was applied to all images in the dataset, to create binary signal masks for each marker. Of note, the algorithm takes into account pixel intensity but also patterns of neighbouring pixels, which could aid the removal of background without losing true signal. Comparison of the images without normalisation, after manual thresholding and the semi-automated background removal, showed that the latter approach could be successfully applied to retain true signal while removing a substantial amount of background noise (Supplementary figure 4). To further confirm the validity of this approach, a t-SNE embedding was computed from the single cell data extracted from the binary signal masks (Figure 3d, supplementary figure 5). Inspection of sample and marker overlays on the t-SNE embedding, using colour-coding, showed cells grouped by marker expression rather than per sample, similar to the manual thresholding (Figure 3c). Furthermore, the signal to noise ratio was higher and cells that were positive for low intensity markers, such as FOXP3 and CD103, were identified by the semi-automated background removal in contrast to manual thresholding (supplementary figure 1, 5).

### Background removal and binarization combined with the proposed downstream analysis pipeline allows phenotyping of the tumour immune microenvironment

To demonstrate the added value of performing background removal and binarization of pixel intensity for the identification of immune-phenotypes, clusters of cells with a comparable marker profile were identified, by applying Gaussian mean shift clustering on the t-SNE embedding, computed in the previous section. Next, clusters were mapped back onto the segmentation masks in ImaCytE. A proliferating and non-proliferating tumour cluster was identified through the expression of keratin and distinguished by Ki-67 (Figure 4a). Five myeloid clusters were identified by their CD68 expression and differentiated by CD204, CD163 and HLA-DR expression. Furthermore, five lymphoid-cell clusters could be identified where two clusters were CD8 positive, thus, corresponding to cytotoxic T cells. Of note, one of these clusters also showed positive signal for keratin indicating that these cells were located on top of epithelial cancer cells. Three clusters were CD8 negative and considered to be mostly composed of CD4 T cells. One of the clusters corresponded to regulatory T cells (FOXP3^+^) and the other two clusters were differentiated by Ki-67 expression. The tumour, myeloid and lymphoid clusters were each mapped back onto the images and compared to the original MCD images in the MCD viewer (Figure 4b). Indeed, the number and location of positive cells for each phenotype was comparable between the images overlaid with phenotype masks and the original MCD images. Thus, background removal using ilastik combined with binarization is applicable to normalise IMC datasets derived from archival samples and allows for the identification and localisation of biologically relevant phenotypes.

**Figure 4.**
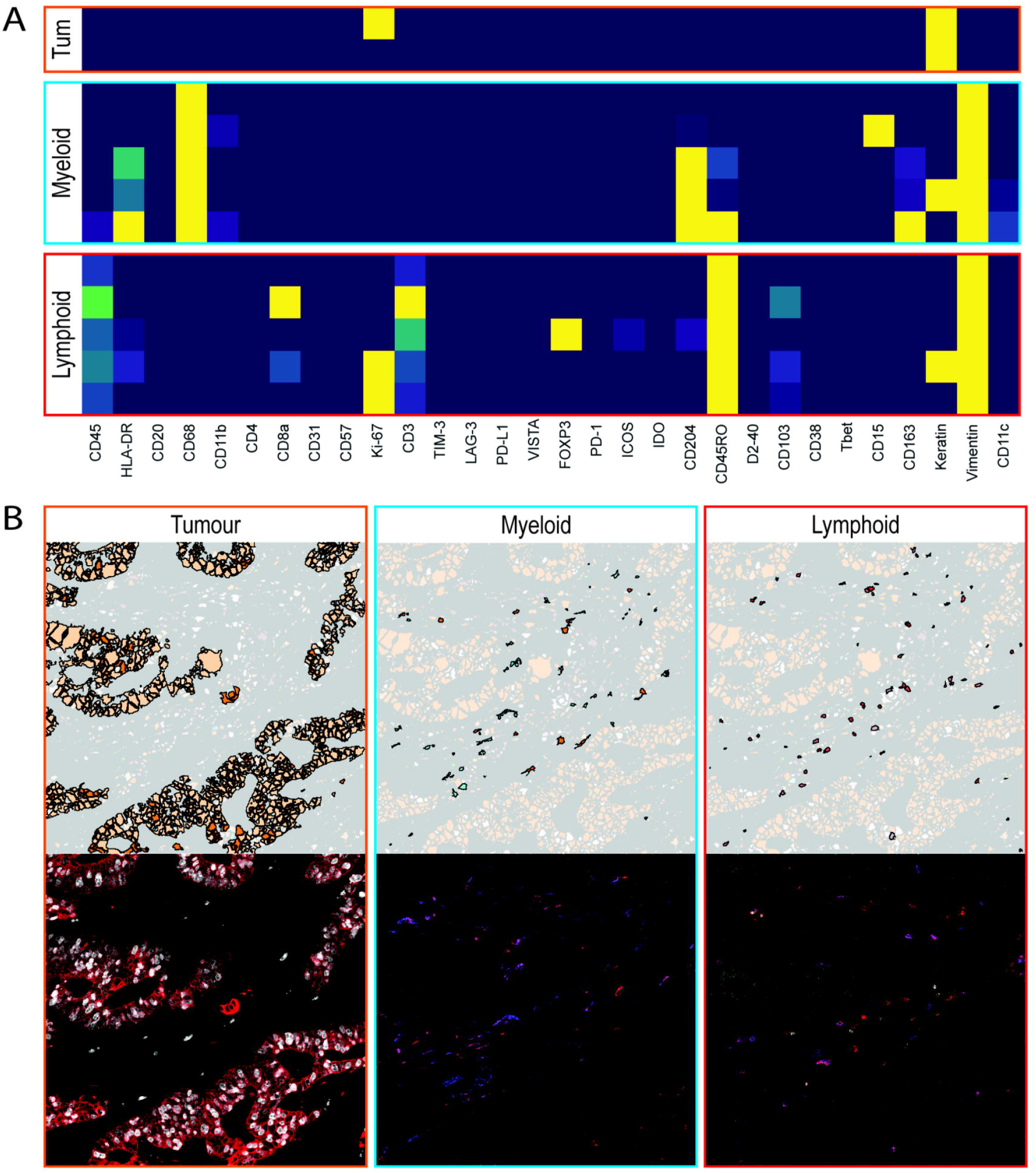
(a) Lymphoid, myeloid and tumour (tum) clusters identified in the t-SNE analysis from figure 3b. (b) the clusters were mapped back onto a representative image and compared to their corresponding cell types in the mcd files. The images contain the following markers: for tumour keratin (red) and Ki67 (white), for myeloid cells CD68 (red) and CD163 (blue) and for lymphoid cells CD3 (red), CD8 (blue) and FOXP3 (cyan). To improve visibility a lower threshold of 1 dual counts was set.

## Discussion

With the rise of mass cytometry for the characterisation of cellular contextures in health and disease, IMC has surfaced as a valuable tool to investigate immunophenotypes while preserving spatial information. IMC allows the simultaneous investigation of over 40 markers thereby generating complex datasets that require analysis tools that combine deep immunophenotyping with spatial localisation and neighbourhood analysis. However, before the interpretation of relevant data, non-biological variation of signal intensities between tissues should be addressed. Technical noise is consistent in each image and will therefore not influence downstream analysis as long as the signal-to-noise ratio is high, and can be addressed by optimising the antibodies and performing the antibody labelling of tissues in a single experiment. However, sample-specific noise, related to tissue processing procedures, is impossible to address during the staining procedures. FFPE tissue is often the tissue of choice for IMC due to its accessibility and good morphology. Differences in ischemia time, tissue fixation and age occur between tissues and this causes the labelling intensity or tissue-derived noise to vary as well, prompting the need to normalise the data before analysis. Direct phenotyping of IMC data, without normalisation, can lead to cells clustering per sample rather than phenotypes as the variation in overall signal intensity overshadows the differences between cell types. Furthermore, the signal range of the more abundant structural markers (e.g. keratin, vimentin) is much wider compared to scarcer, but important targets of investigation (e.g. co-receptors on T cells). This difference influences downstream computational methods such as clustering or dimensionality reduction algorithms and hinders the detection of specific cell subsets.

In this work, we enable the analysis of large IMC datasets and improve the detection of cells expressing lowly abundant proteins by removing background noise and binarizing each sample’s pixel values. Binarizing was done by assigning the value of 1 to the pixels that express a marker, regardless of the sample or markers noise and the value of 0 to all the other pixels. More specifically, we tested two different normalisation approaches. First, a manual method was utilised where a lower threshold was set for each marker in all images. All pixels below threshold were set to 0 and all pixels above were set to 1. Then the marker expression per cell was defined as the mean of all pixels in a cell. This approach indeed overcame variation between tissues and allowed for t-SNE-guided phenotype identification. However, manual thresholding is labour intensive and relies on vast knowledge of the expected labelling patterns for each antibody, leading to potentially biased results. In general, thresholding is a trade-off between the removal of background and loss of true signal and therefore can result in the frequent designation of false negatives. An automated approach could overcome these challenges but large variations between samples and markers make fully automated unsupervised methods infeasible and the lack of labelled datasets is prohibitive for supervised machine learning approaches. Therefore, we proposed a, semi-automated, methodology using ilastik where we first annotate representative pixels either as actual signal or background noise and then a random forest classifier is run to categorise the whole dataset, based on these categories. This approach is faster, less subjective and results in data comparable to manual thresholding. Furthermore, loss of true signal is less frequently observed, and dim markers are more clearly reflected after semi-automated background removal and pixel intensity binarization.

In recent years, great advancements have been made on the analysis and interpretation of single cell mass cytometry data and corresponding tools also prove invaluable for the analysis of IMC data. However, these should be combined with the knowledge of the immunohistochemistry field on tissue specific artefacts and variations, such as intensity differences and sticking of antibodies to necrotic regions. Recent publications have improved the labelling and imaging methodologies and simultaneously new tools for the interpretation of IMC data have emerged. As IMC becomes more accessible, many studies will utilise FFPE tissues of different subjects, and will thus have to overcome the tissue specific variations this brings. The here described normalisation methodology enables the analysis of datasets consisting of different tissues and the easier detection of lowly abundant proteins allowing the use of IMC phenotyping and interpretation tools. Furthermore, the methodology does not call for changes in the antibody labelling procedure but can be directly applied on previously obtained datasets. Thus, this work has the potential to directly aid research groups in their analysis and interpretation of imaging mass cytometry data.

## Data availability statement

Data is available upon request.

## References

1. de Vries NL, van Unen V, Ijsselsteijn ME, Abdelaal T, van der Breggen R, Farina Sarasqueta A, et al. High-dimensional cytometric analysis of colorectal cancer reveals novel mediators of antitumour immunity. Gut. 2020;69(4):691–703.

2. Rubin SJS, Bai L, Haileselassie Y, Garay G, Yun C, Becker L, et al. Mass cytometry reveals systemic and local immune signatures that distinguish inflammatory bowel diseases. Nat Commun. 2019;10(1):2686.

3. Li N, van Unen V, Abdelaal T, Guo N, Kasatskaya SA, Ladell K, et al. Memory CD4(+) T cells are generated in the human fetal intestine. Nat Immunol. 2019;20(3):301–12.

4. Bendall SC, Simonds EF, Qiu P, Amir el AD, Krutzik PO, Finck R, et al. Single-cell mass cytometry of differential immune and drug responses across a human hematopoietic continuum. Science. 2011;332(6030):687–96.

5. Newell EW, Sigal N, Bendall SC, Nolan GP, Davis MM. Cytometry by time-of-flight shows combinatorial cytokine expression and virus-specific cell niches within a continuum of CD8+ T cell phenotypes. Immunity. 2012;36(1):142–52.

6. Wei SC, Levine JH, Cogdill AP, Zhao Y, Anang NAS, Andrews MC, et al. Distinct Cellular Mechanisms Underlie Anti-CTLA-4 and Anti-PD-1 Checkpoint Blockade. Cell. 2017;170(6):1120–33 e17.

7. Bandura DR, Baranov VI, Ornatsky OI, Antonov A, Kinach R, Lou X, et al. Mass cytometry: technique for real time single cell multitarget immunoassay based on inductively coupled plasma time-of-flight mass spectrometry. Anal Chem. 2009;81(16):6813–22.

8. Ornatsky OI, Kinach R, Bandura DR, Lou X, Tanner SD, Baranov VI, et al. Development of analytical methods for multiplex bio-assay with inductively coupled plasma mass spectrometry. J Anal At Spectrom. 2008;23(4):463–9.

9. Giesen C, Wang HA, Schapiro D, Zivanovic N, Jacobs A, Hattendorf B, et al. Highly multiplexed imaging of tumor tissues with subcellular resolution by mass cytometry. Nat Methods. 2014;11(4):417–22.

10. Chevrier S, Crowell HL, Zanotelli VRT, Engler S, Robinson MD, Bodenmiller B. Compensation of Signal Spillover in Suspension and Imaging Mass Cytometry. Cell Syst. 2018;6(5):612–20 e5.

11. Ijsselsteijn ME, van der Breggen R, Farina Sarasqueta A, Koning F, de Miranda N. A 40-Marker Panel for High Dimensional Characterization of Cancer Immune Microenvironments by Imaging Mass Cytometry. Front Immunol. 2019;10:2534.

12. Guo N, van Unen V, Ijsselsteijn ME, Ouboter LF, van der Meulen AE, Chuva de Sousa Lopes SM, et al. A 34-Marker Panel for Imaging Mass Cytometric Analysis of Human Snap-Frozen Tissue. Front Immunol. 2020;11:1466.

13. Schulz D, Zanotelli VRT, Fischer JR, Schapiro D, Engler S, Lun XK, et al. Simultaneous Multiplexed Imaging of mRNA and Proteins with Subcellular Resolution in Breast Cancer Tissue Samples by Mass Cytometry. Cell Syst. 2018;6(4):531.

14. Jackson HW, Fischer JR, Zanotelli VRT, Ali HR, Mechera R, Soysal SD, et al. The single-cell pathology landscape of breast cancer. Nature. 2020;578(7796):615–20.

15. Wang YJ, Traum D, Schug J, Gao L, Liu C, Consortium H, et al. Multiplexed In Situ Imaging Mass Cytometry Analysis of the Human Endocrine Pancreas and Immune System in Type 1 Diabetes. Cell Metab. 2019;29(3):769–83 e4.

16. Damond N, Engler S, Zanotelli VRT, Schapiro D, Wasserfall CH, Kusmartseva I, et al. A Map of Human Type 1 Diabetes Progression by Imaging Mass Cytometry. Cell Metab. 2019;29(3):755–68 e5.

17. Berg S, Kutra D, Kroeger T, Straehle CN, Kausler BX, Haubold C, et al. ilastik: interactive machine learning for (bio)image analysis. Nat Methods. 2019;16(12):1226–32.

18. Carpenter AE, Jones TR, Lamprecht MR, Clarke C, Kang IH, Friman O, et al. CellProfiler: image analysis software for identifying and quantifying cell phenotypes. Genome Biol. 2006;7(10):R100.

19. Schapiro D, Jackson HW, Raghuraman S, Fischer JR, Zanotelli VRT, Schulz D, et al. histoCAT: analysis of cell phenotypes and interactions in multiplex image cytometry data. Nat Methods. 2017;14(9):873–6.

20. Somarakis A, Van Unen V, Koning F, Lelieveldt BPF, Hollt T. ImaCytE: Visual Exploration of Cellular Microenvironments for Imaging Mass Cytometry Data. IEEE Trans Vis Comput Graph. 2019.

21. Economou M, Schoni L, Hammer C, Galvan JA, Mueller DE, Zlobec I. Proper paraffin slide storage is crucial for translational research projects involving immunohistochemistry stains. Clin Transl Med. 2014;3(1):4.

22. Werner M, Chott A, Fabiano A, Battifora H. Effect of formalin tissue fixation and processing on immunohistochemistry. Am J Surg Pathol. 2000;24(7):1016–9.

23. O’Hurley G, Sjostedt E, Rahman A, Li B, Kampf C, Ponten F, et al. Garbage in, garbage out: a critical evaluation of strategies used for validation of immunohistochemical biomarkers. Mol Oncol. 2014;8(4):783–98.

24. Bussolati G, Leonardo E. Technical pitfalls potentially affecting diagnoses in immunohistochemistry. J Clin Pathol. 2008;61(11):1184–92.

25. Taylor CR, Levenson RM. Quantification of immunohistochemistry--issues concerning methods, utility and semiquantitative assessment II. Histopathology. 2006;49(4):411–24.

26. Leong AS. Quantitation in immunohistology: fact or fiction? A discussion of variables that influence results. Appl Immunohistochem Mol Morphol. 2004;12(1):1–7.

27. van Unen V, Hollt T, Pezzotti N, Li N, Reinders MJT, Eisemann E, et al. Visual analysis of mass cytometry data by hierarchical stochastic neighbour embedding reveals rare cell types. Nat Commun. 2017;8(1):1740.

28. Höllt T, Pezzotti N, van Unen V, Koning F, Eisemann E, Lelieveldt B, et al. Cytosplore: Interactive Immune Cell Phenotyping for Large Single-Cell Datasets. Computer Graphics Forum. 2016;35(3):171–80.

