## Supplementary Table and Figures for "Semi-automated background removal limits loss of data and normalises the images for downstream analysis of imaging mass cytometry data"

**Supplementary figures**

Supplementary table 1| Imaging mass cytometry antibody panel. Antibodies with a * had prediluted stock solutions

| Target | Clone | Metal | Incubation Time | Temperature | Dilution |
|---|---|---|---|---|---|
| CD45 | D9M8I | $^{115}In$ | Overnight | 4°C | 50 |
| HLA-DR | TAL-1B5 | $^{141}Pr$ | 5 hours | RT | 100* |
| CD20 | H1 | $^{142}Nd$ | Overnight | 4°C | 100 |
| CD68 | D4B9C | $^{143}Nd$ | Overnight | 4°C | 100* |
| CD11b | D6X1N | $^{144}Nd$ | 5 hours | RT | 100 |
| CD4 | EPR6855 | $^{145}Nd$ | 5 hours | RT | 50 |
| CD8α | D8A8Y | $^{146}Nd$ | 5 hours | RT | 50 |
| CD31 | 89C2 | $^{147}Sm$ | Overnight | 4°C | 100* |
| CD57 | HNK-1/Leu-7 | $^{151}Eu$ | Overnight | 4°C | 100* |
| Ki-67 | 8D5 | $^{152}Sm$ | Overnight | 4°C | 100* |
| CD3 | D7A6E | $^{153}Eu$ | Overnight | 4°C | 50 |
| TIM-3 | D5D5R | $^{154}Sm$ | 5 hours | RT | 100 |
| LAG3 | D2G4O | $^{155}Gd$ | 5 hours | RT | 50 |
| PD-L1 | E1L3N | $^{156}Gd$ | Overnight | 4°C | 50 |
| VISTA | D1L2G | $^{158}Gd$ | 5 hours | RT | 100 |
| FoxP3 | D6O8R | $^{159}Tb$ | Overnight | 4°C | 50 |
| PD-1 | D4W2J | $^{160}Gd$ | 5 hours | RT | 50 |
| ICOS | D1K2T | $^{161}Dy$ | 5 hours | RT | 50 |
| IDO | D5J4E | $^{162}Dy$ | Overnight | 4°C | 100 |
| CD204 | J5HTR3 | $^{164}Dy$ | 5 hours | RT | 50 |
| CD45RO | UCHL1 | $^{165}Ho$ | Overnight | 4°C | 100* |
| D2-40 | D2-40 | $^{166}Er$ | Overnight | 4°C | 100* |
| CD103 | EPR4166(2) | $^{168}Er$ | 5 hours | RT | 50 |
| CD38 | EPR4106 | $^{169}Tm$ | Overnight | 4°C | 100* |
| T-bet | 4B10 | $^{170}Er$ | 5 hours | RT | 50 |
| CD15 | BRA-4F1 | $^{171}Yb$ | Overnight | 4°C | 100* |
| CD163 | EPR14643-36 | $^{173}Yb$ | 5 hours | RT | 50 |
| CD11c | EP1347Y | $^{176}Yb$ | 5 hours | RT | 100 |
| Vimentin | D21H3 | $^{194}Pt$ | Overnight | 4°C | 50 |
| Pan-Keratin | AE1/AE3 and C11 | $^{198}Pt$ | Overnight | 4°C | 50 |

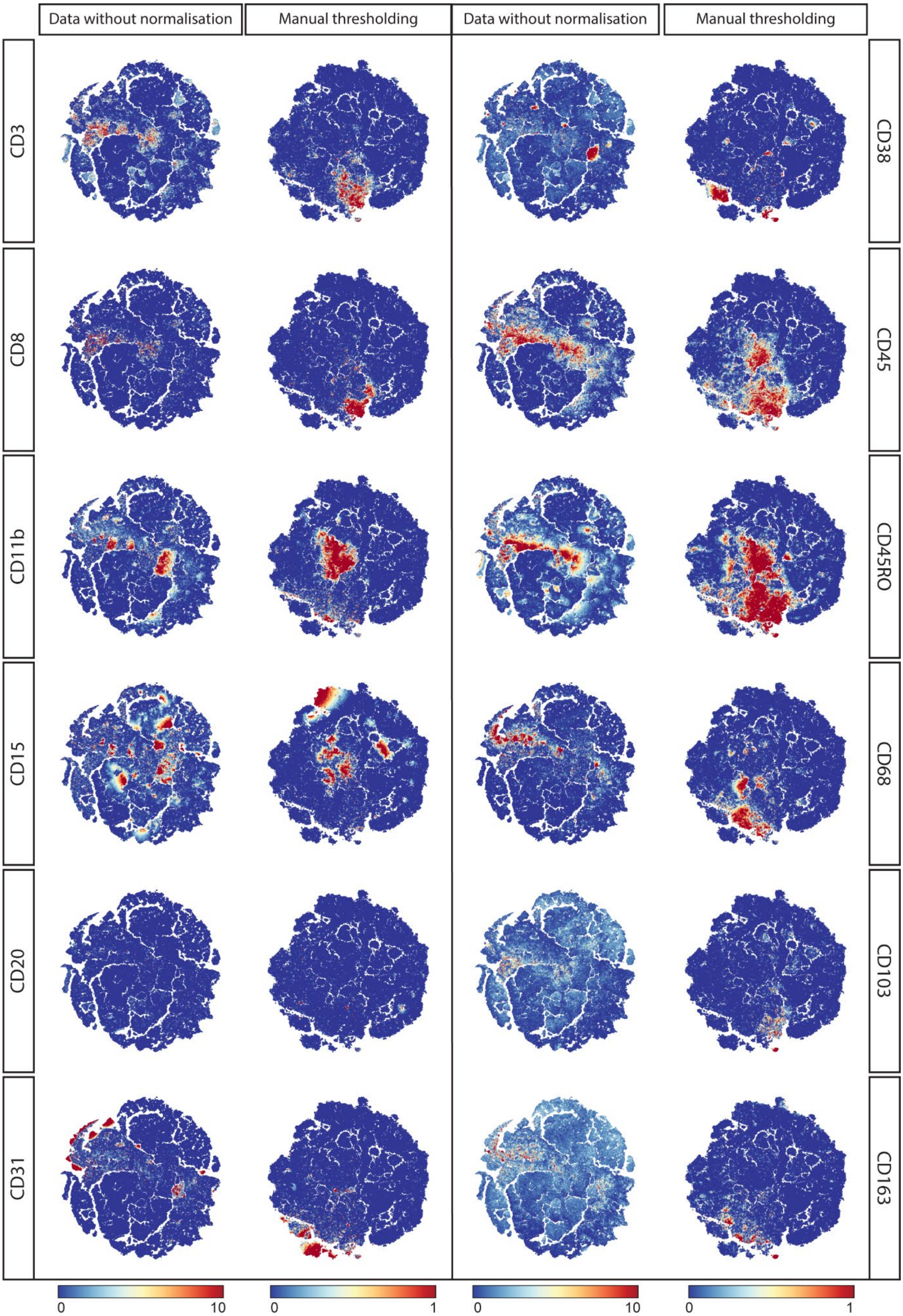
Data without normalisation
Manual thresholding
Data without normalisation
Manual thresholding
CD3
CD8
CD11b
CD15
CD20
CD31
CD38
CD45
CD45RO
CD68
CD103
CD163
0
10
0
1
0
10
0
1

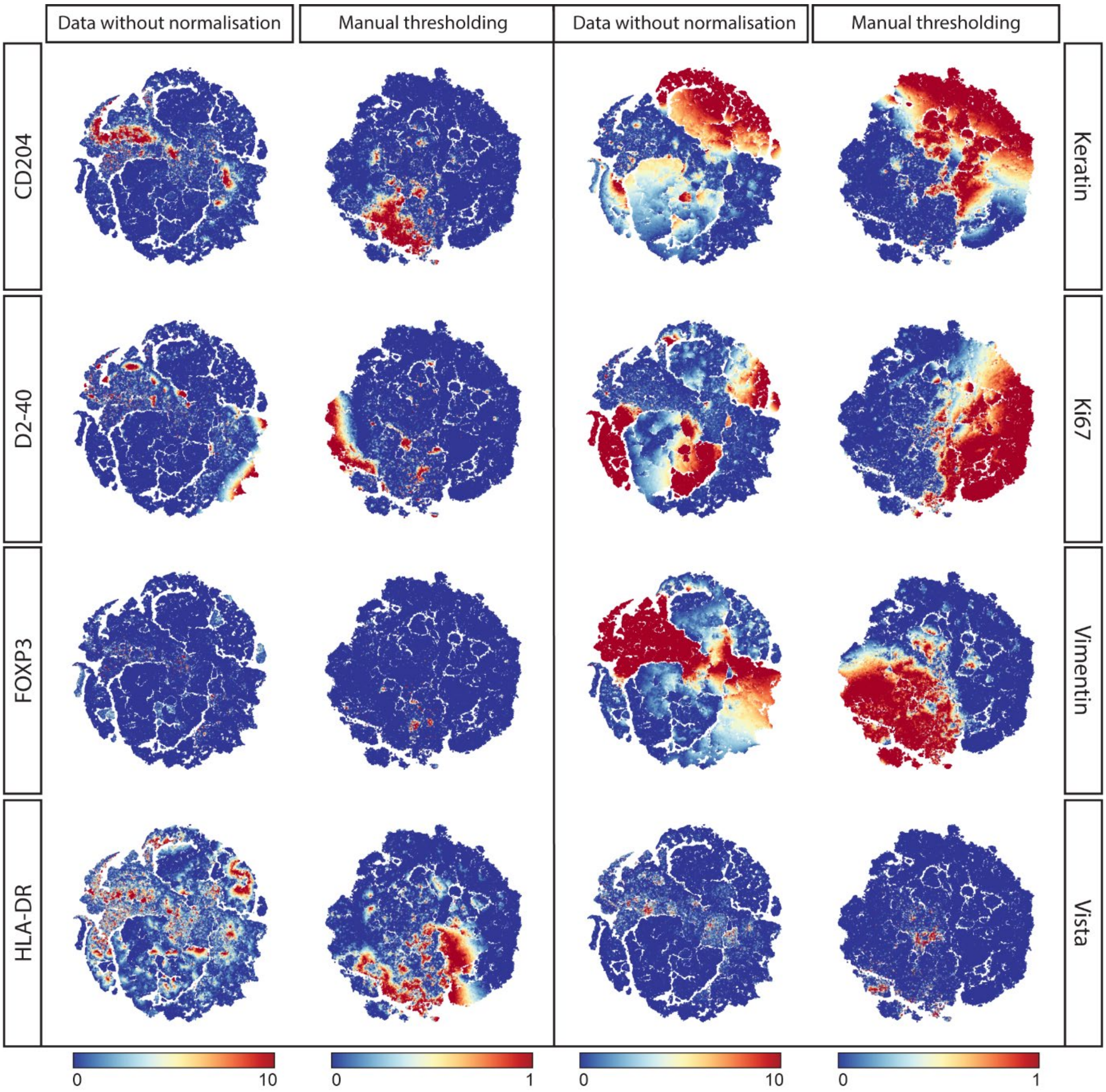


Supplementary figure 1| TSNE analysis on single cells extracted from images without normalisation compared to manually thresholded & binarized images.

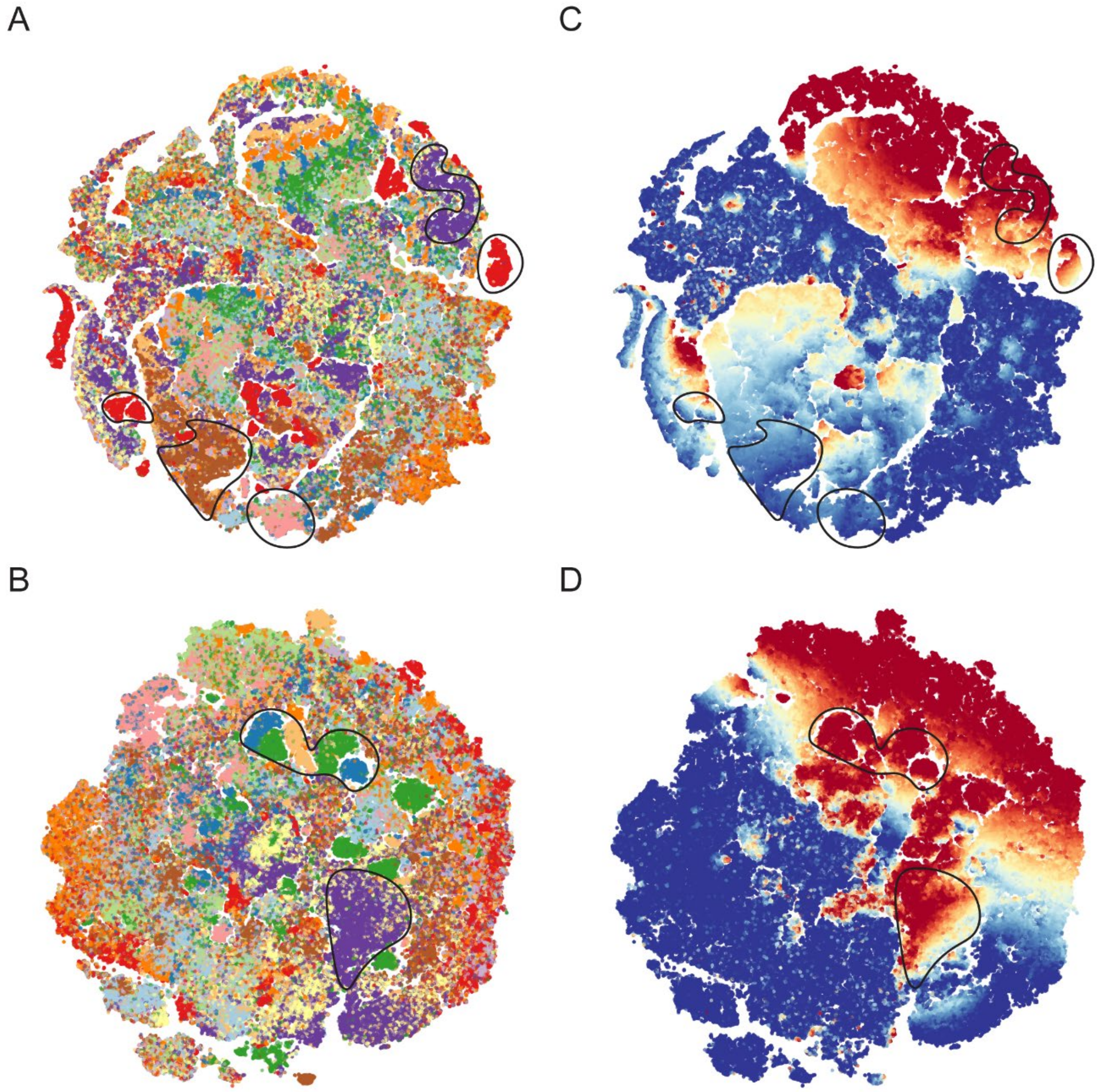


Supplementary figure 2| (a) TSNE embedding of single cell data extracted from images without normalisation. Each colour represents a different sample and cells cluster together by phenotype. Sample-specific clustering is encircled. (b) TSNE embedding of single cell data extracted from images that underwent manual thresholding. Each colour represents a different sample and sample specific clustering is encircled. (c and d) Keratin expression overlay on the TSNE from non-normalised data (c) and data that underwent manual thresholding (d). sample specific clusters were highlighted

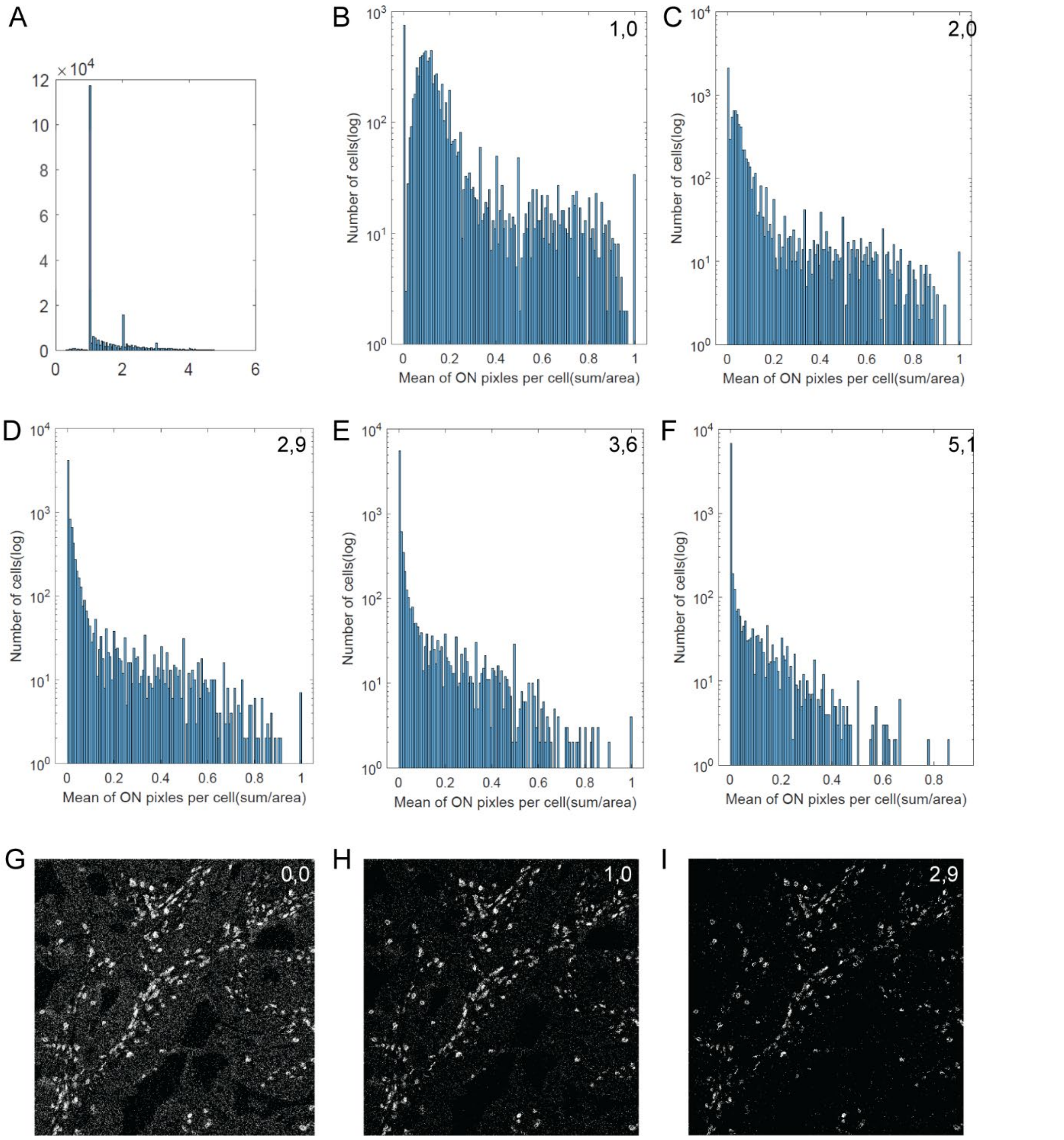


Supplementary figure 3| (a) pixel data plotted in a histogram with on the x-axis the pixel intensity and on the y-axis the frequency. (b) CD3 expression in one image shown as the mean positive pixels per cell with a cut off threshold at 1. ). All pixels below threshold were set to 0 and above threshold at 1 and used to determine the mean intensity per cell. (c-f) histograms of CD3 expression with different threshold between 2 and 5. (g-i) CD3 expression in the image corresponding to a threshold of 0, 1 or 2,9.

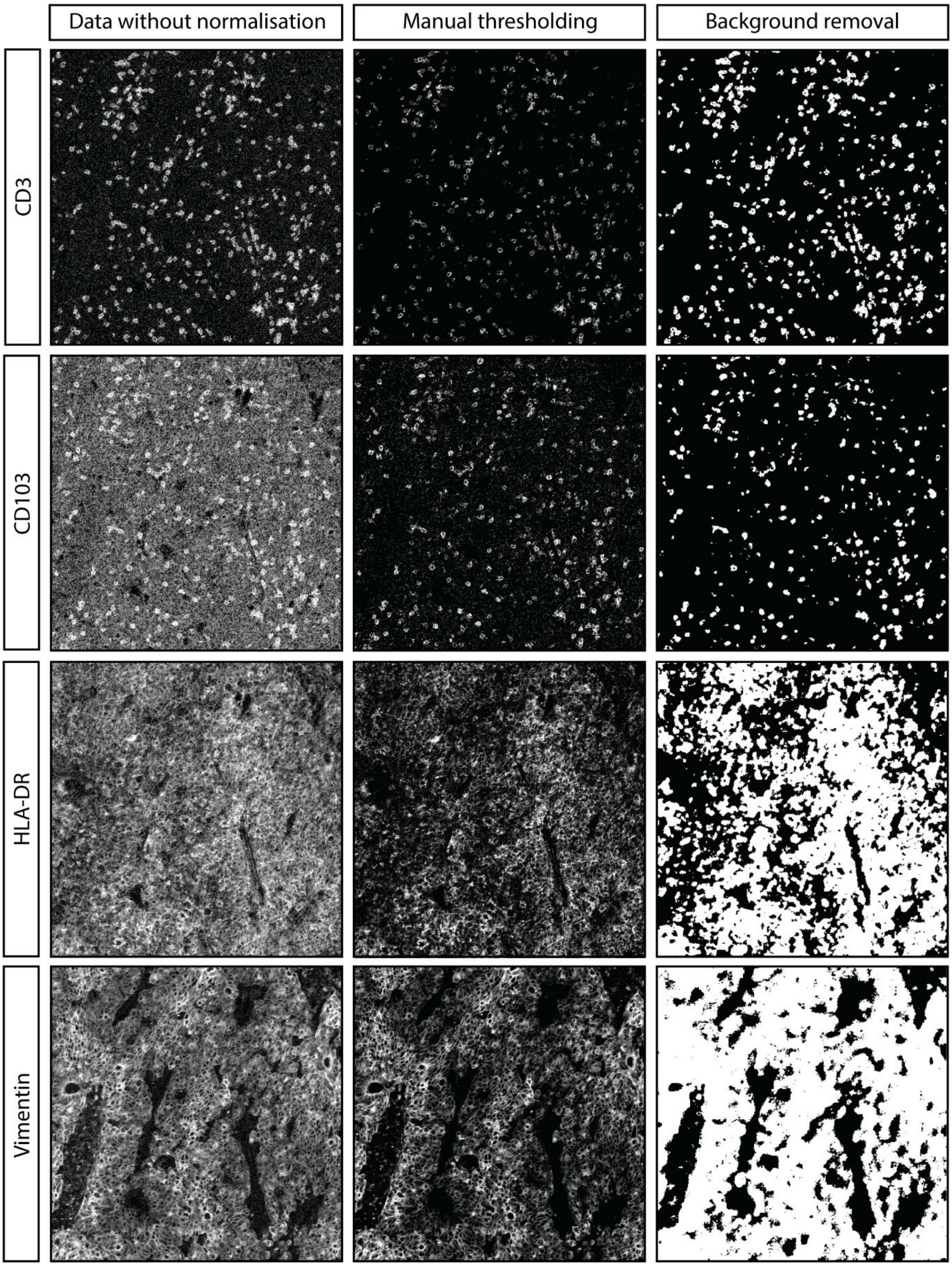


Supplementary figure 4| Comparison of antibody signal between images without normalisation, after manual thresholding and after background removal

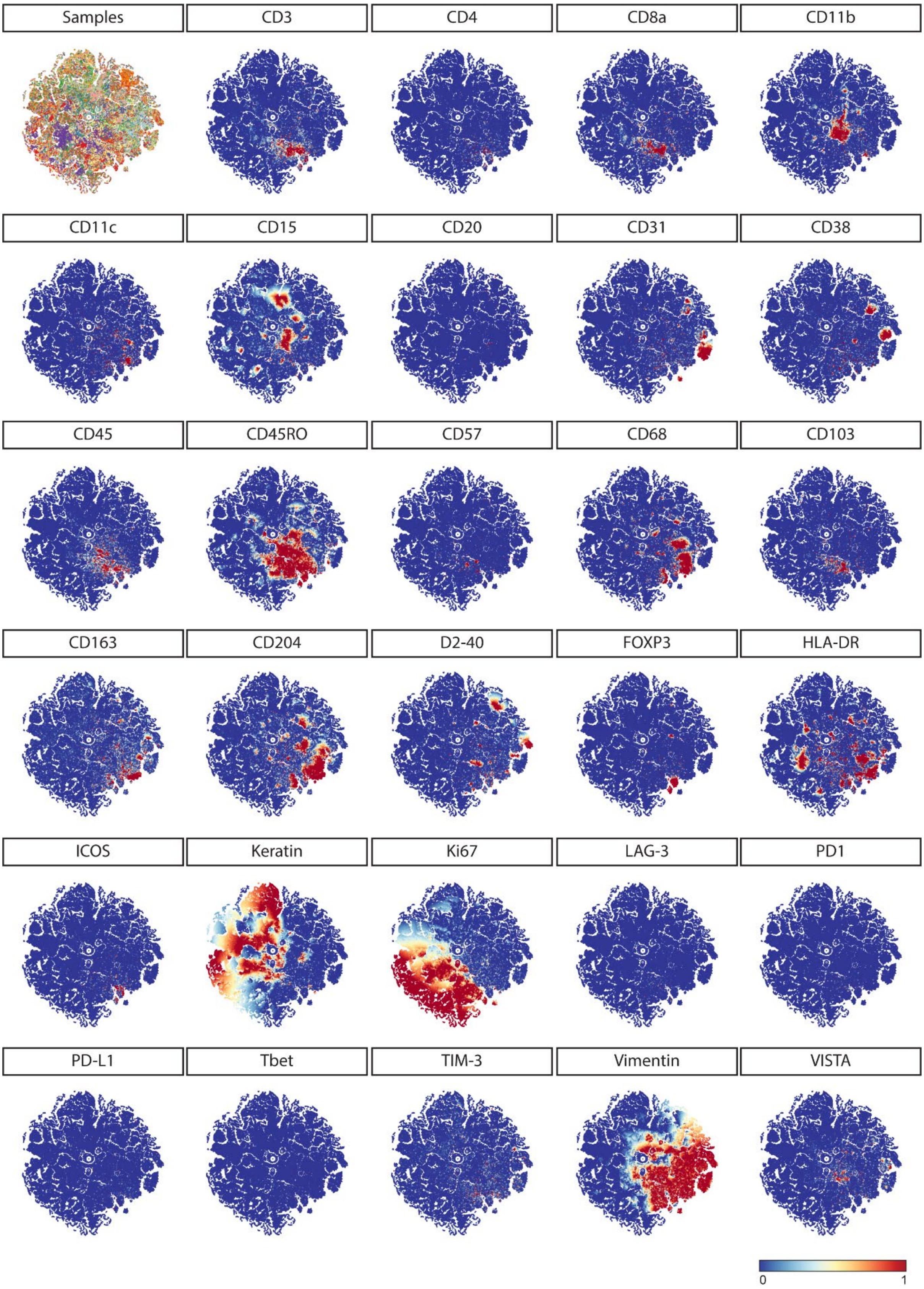


Supplementary figure 5| TSNE analysis on single cells extracted from binary images created by background removal in Ilastik. Data is shown in a range of 0 (blue) to 1 (red).
